# Kappa opioid receptor modulation on GABAergic inputs onto ventral periaqueductal gray dopamine neurons

**DOI:** 10.1101/389536

**Authors:** Chia Li, Thomas L. Kash

## Abstract

The kappa opioid receptor (KOR) system has been implicated in regulation of many behaviors including pain. While there are numerous studies suggesting KOR-regulation of pain being mediated spinally, there have been reports of pain-like behaviors regulated by central KOR signaling. In particular, oxytocin-induced analgesia appears to be mediated by KOR receptors within the ventrolateral periaqueductal gray (vlPAG). The vlPAG is a brain region that has long been known to be involved in the regulation of pain. We recently found that dopamine (DA) neurons within the vlPAG represent a specific population of neurons that can regulate pain-like behaviors. In this study, we sought to determine the impact of KOR signaling on GABAergic inputs to the vlPAG DA neurons, and determine the mechanism of inhibition. We found that activation of KOR significantly reduced GABAergic transmission onto vlPAG DA neurons. In addition, our data suggest this effect is mediated pre-synaptically via the G-protein βγ subunit. These data suggest the possibility that KOR-activation disinhibits vlPAG dopamine neurons, which could lead to altered regulation of pain-related behaviors.

## Introduction

As a member of the opioid receptor family, kappa opioid receptor actions have been shown to contribute important roles in analgesia, mood modulation, and drug abuse [1– 4]. They are widely spread throughout the brain, spinal cord, and peripheral systems [5], and are activated endogenously via the opioid peptide dynorphin [6]. Canonically, KOR-activation involves the inhibition of cyclic AMP production [7] that is mediated through the coupling of the inhibitory G-protein Gi/o [8]. Through the dissociation of the α subunit, which in turns recruit beta arrestin and activate downstream mitogen-activated protein (MAP) kinases that affect transcription factor expression, such as ERK1/2 [9] and p38. The βγ subunit can directly bind and inhibit calcium channels, as well as increase potassium channel conductance [10]. These direct effects on ion channel conductance have been found in several brain regions, ranging from the hippocampus to the dorsal root ganglia [11]. In addition, the phosphorylated KOR activates ERK1/2, as well as phosphoinositide 3 (PI_3_) kinase and protein kinase A (PKA). Evidence suggests ERK signaling mediates KOR activation-induced attenuation in inhibitor transmission in the BNST [12]. Further, studies have shown that p38 MAP kinase signaling can regulate KOR-mediated inhibition of glutamate transmission in the BNST, and is required for negative affective behaviors that can be blocked by KOR antagonists [13]. The divergence in signaling pathways that mediate the effects of KOR activation are meaningful, as it suggests that biased agonists could be designed to selectively target specific pathways to engender different effects.

The A10dc group DA neurons project from the ventral lateral periaqueductal gray (vlPAG) to the extended amygdala - the bed nucleus of stria terminalis (BNST) and the central amygdala (CeA), areas known to regulate stress, anxiety and pain-related behaviors [14–17]. Recent studies from our group have found that chemogenetic activation of these vlPAG DA neurons can alter pain-related behaviors. Beyond pain, the fucntion of these neurons has been implicated in arousal [18] and social behavior [19]. Of note, anti-nociception induced by oxytocin can be blocked by Kappa Opioid Receptor (KOR) antagonism in the PAG [20]. Mechanistically, it has been shown that kappa opioid receptor activation can modulate dopaminergic neurons in the VTA. More specifically, KOR agonist inhibits the VTA DA neurons projecting to the prefrontal cortex [21]. While isolating GABA-mediated inhibitory post-synaptic currents (IPSCs) in the VTA dopamine neurons, the same research group found that opioid agonists inhibited GABAergic transmission. Additionally, GABAergic neurons have been shown to inhibit projection dopamine neurons in the VTA [22]. Similar to the VTA, a population of tonically-active GABAergic neurons have been found in the PAG [23]. Although KORs have been shown to distribute widely in the PAG [24], specific dopamine neuron-modulation has yet to be probed due to the heterogeneity of the PAG. The existing behavioral and electrophysiological findings together lead us to the hypothesis that kappa opioid receptors could serve a role in modulating GABAergic inputs onto the vlPAG dopaminergic neurons.

## Materials and Methods

### Animals and Husbandry

Male TH-eGFP mice on a Swiss Webster background (aged between 5 to 9 weeks) were bred and used in accordance with an animal use protocol approved by the University of North Carolina – Chapel Hill (IACUC). Mice were group-housed in our colony room under a 12:12-hour light cycle, with lights on at 7:00 AM daily. Mice were given *ad libitum* access to rodent chow and water. Mating pairs of mice were created by GENSAT and obtained from the Mutant Mouse Regional Resource Center in North Carolina. In the TH-eGFP mouse line, the genome was modified to contain multiple copies of a modified BAC in which an eGFP reporter gene was inserted immediately upstream of the coding sequence of the gene for tyrosine hydroxylase (TH). Data presented here were obtained from the transgenic mice maintained in-house.

### Electrophysiology Brain Slice Preparation

Mice were decapitated under isoflurane anesthesia and their brains were rapidly removed and placed in ice-cold sucrose artificial cerebrospinal fluid (ACSF): (in mM) 194 sucrose, 20 NaCl, 4.4 KCl, 2 CaCl2, 1 MgCl2, 1.2 NaH2PO4, 10.0 glucose, and 26.0 NaHCO3 saturated with 95% O2/5% CO2. Three hundred micron slices were prepared using a Leica VT1200 vibratome (Wetzlar, Germany).

### Slice Whole-Cell Electrophysiology

Brain slices containing PAG were obtained and stored at approximately 30°C in a heated, oxygenated holding chamber containing artificial cerebrospinal fluid (ACSF) (in mmol/L) 124NaCl, 4.4 KCl, 2 CaCl2, 1.2 MgSO4, 1 NaH2PO4, 10.0 glucose, and 26.0 sodium bicarbonate before being transferred to a submerged recording chamber maintained at approximately 30°C (Warner Instruments, Hamden, Connecticut) Recording electrodes (3–5 MΩ) were pulled with a Flaming-Brown Micropipette Puller (Sutter Instruments, Novato, CA), using thin-walled borosilicate glass capillaries. During inhibitory transmission experiments, recording electrodes were filled with (in mmol/L) 70 KCl, 65 K+-gluconate, 5 NaCl, 10 4-(2-hydroxyethyl)-1-piperazineethanesulfonic acid, QX-314, .6 EGTA, 4 ATP, .4 GTP, pH 7.4, 290 to 295 mOsmol. In experiments where post-synaptic GPCR signaling was blocked, GDPβs was used to replace GTP in the internal solution. All experiments were conducted under the voltage clamp configuration, cells were held at −70 mV and inhibitory post-synaptic currents (IPSCs) were pharmacologically isolated with 3 mmol/L kynurenic acid, to block α-amino-3-hydroxy-5-methyl-4-isoxazole-propionic acid (AMPA) and *N*-methyl-D-aspartate (NMDA) receptor-dependent post-synaptic current. To isolate miniature inhibitory post-synaptic currents (mIPSCs), tetrodotoxin (0.5 μmol/L) was added to the perfusing ACSF solutions described above. Signals were acquired via a Multiclamp 700B amplifier (Molecular Devices, Sunnyvale, California), digitized at 20 kHz, filtered at 3 kHz, and analyzed using Clampfit 10.2 software (Molecular Devices). Input resistance and access resistance were continuously monitored during experiments. Experiments in which changes in access resistance were greater than 20% were not included in the data analysis.

### Statistics

Effects of drugs during electrophysiological recordings were evaluated by comparing the magnitude of the dependent measure (mIPSC frequency and amplitude) between the baseline and wash-on (when drug had reached maximal effect at 10 minutes) periods using paired *t* tests. The effects of antagonists/blockers on the ability of drugs to modulate synaptic transmission were compared using *t* tests during the washout period. All values given for drug effects throughout the article are presented as mean ± SEM.

### Drugs

Dynorphin A (300 nM), Norbinaltorphimine (Nor-BNI, 100 nM) were purchased from Tocris (Ellisville, MO) and dissolved in distilled water. BAPTA-AM (50µM), Gallein (100µM), and Wortmannin (1µM) were purchased from Tocris and dissolved in DMSO. 4-[4-(4-Fluorophenyl)-2-[4-(methylsulfinyl)phenyl]-1H-imidazol-5-yl]pyridine (SB203580, 20 µM) and 4-aminopyridine (4-AP, 100µM) were from purchased from Ascent and dissolved in distilled water; alpha-[Amino[(4-aminophenyl)thio]methylene]-2-(trifluoromethyl)benzeneacetonitrile (SL327, 10 µM) was from Ascent and dissolved in DMSO. Tetrodotoxin citrate (TTX, 500nM), kynurenic acid (3mM), and CTOP (1µM) were purchased from Abcam and dissolved in water. GDPβs (4mM) and RP-Adenosine 3’,5’-cyclic monophophorothioate triethylammonium salt hydrate (RP-camps, 10µM) were purchased from Sigma-Aldrich and dissolved in water. EGTA (100µM) was obtained from Fisher Scienctific and dissolved in 1M NaOH.

## Results

### Endogenous KOR agonist Dynorphin A attenuates GABAergic input onto vlPAG DA neurons via presynaptic mechanisms

We first examined the effects of KOR activation on GABA synaptic transmission via bath application of the endogenous ligand dynorphin A (300nM). A 10-minute bath application of dynorphin A significantly attenuated the mini inhibitory post-synaptic current (mIPSC) in the vlPAG dopamine neurons (Fig1A, n=5). Specifically, a decrease was seen in mIPSC frequency (69.5±7.4% of baseline, p=0.03, Fig1B,D), but not amplitude (99.3±4.8% of baseline, Fig1C,E), suggesting a pre-synaptic mechanism. To further confirm that the dynorphin effect observed was mediated through KOR activation, we incubated the slices in a selective KOR antagonist, nor-BNI (100nM) for 40 min before and during dynorphin wash-on (n=5). In the presence of nor-BNI, dynorphin A application failed to produce effects on either mIPSC frequency (93.3±10.3% of baseline, Fig1F) or amplitude (89.4±6.1% of baseline, Fig1G). To assess the level of tonic KOR functions, we investigated the effects of nor-BNI alone (n=6) and found no effects in sIPSC frequency (100.3±5.2% of baseline, Fig1H) and amplitude (84.8±7.7% of baseline, Fig1I), indicating no tonic KOR activation in the vlPAG DA neurons. These data suggest a pre-synaptic effect of dynorphin in the vlPAG DA neurons. We further verified this pre-synaptic mechanism via blockage of post-synaptic GPCR functions by the replacement of GTP with GDPβs in the recording pipette, disrupting the exchange of GTP and GDP, thus interfering with downstream signaling cascade upon GPCR activation. With post-synaptic GPCR functions impaired, the application of dynorphin A still decreased mIPSC frequency (54.7±5.8% of baseline, n=6, p=0.04, Fig2A,C), but not amplitude (97.8±11.3% of baseline, Fig2B, D), providing additional support that dynorphin attenuates inhibitory input onto vlPAG DA neurons via a pre-synaptic mechanism.

**Figure 1.**
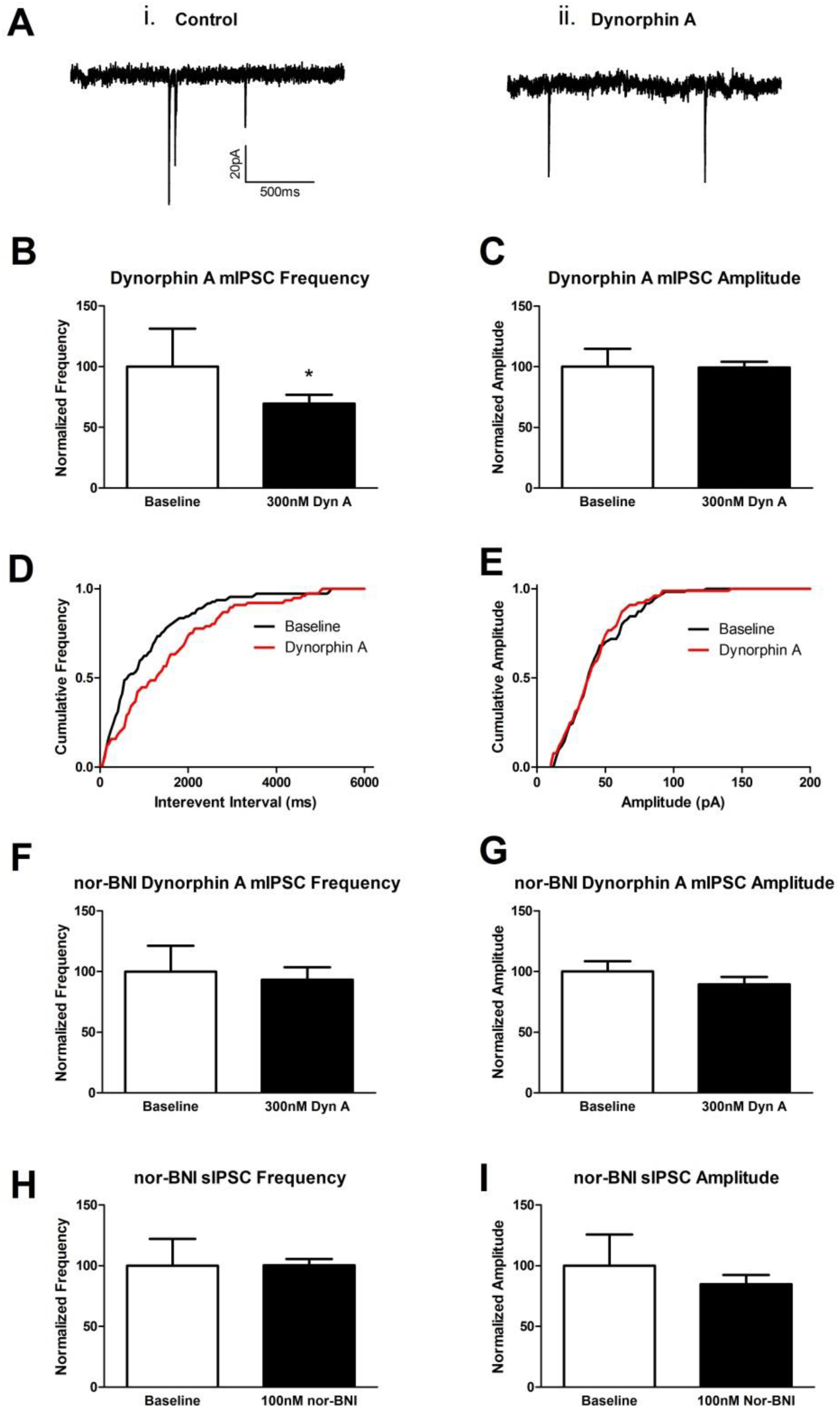
KOR agonist, dynorphin A (300nM), decreases mIPSCs in vlPAG DA neurons via KOR actions. (A) Representative mIPSC traces of baseline control (i) and after 10 minutes of 300nM dynorphin A wash-on (ii). (B) Dynorphin A significantly decreased mIPSC frequency. (C) Dynorphin A had no effect on mIPSC amplitude. (D) Cumulative frequency of mIPSC was shifted towards longer interevent intervals by Dynorphin A. (E) Cumulative probability of mIPSC amplitude was not affected by dynrophin A. (F) Dynorphin A no longer decreased mIPSC frequency in the presence of nor-BNI (100nM). (G) Dynorphin A had no effect on mIPSC amplitude in the presence of nor-BNI. (H) nor-BNI alone had no effect on mIPSC frequency. (I) nor-BNI alone had no effect on mIPSC amplitude

**Figure 2.**
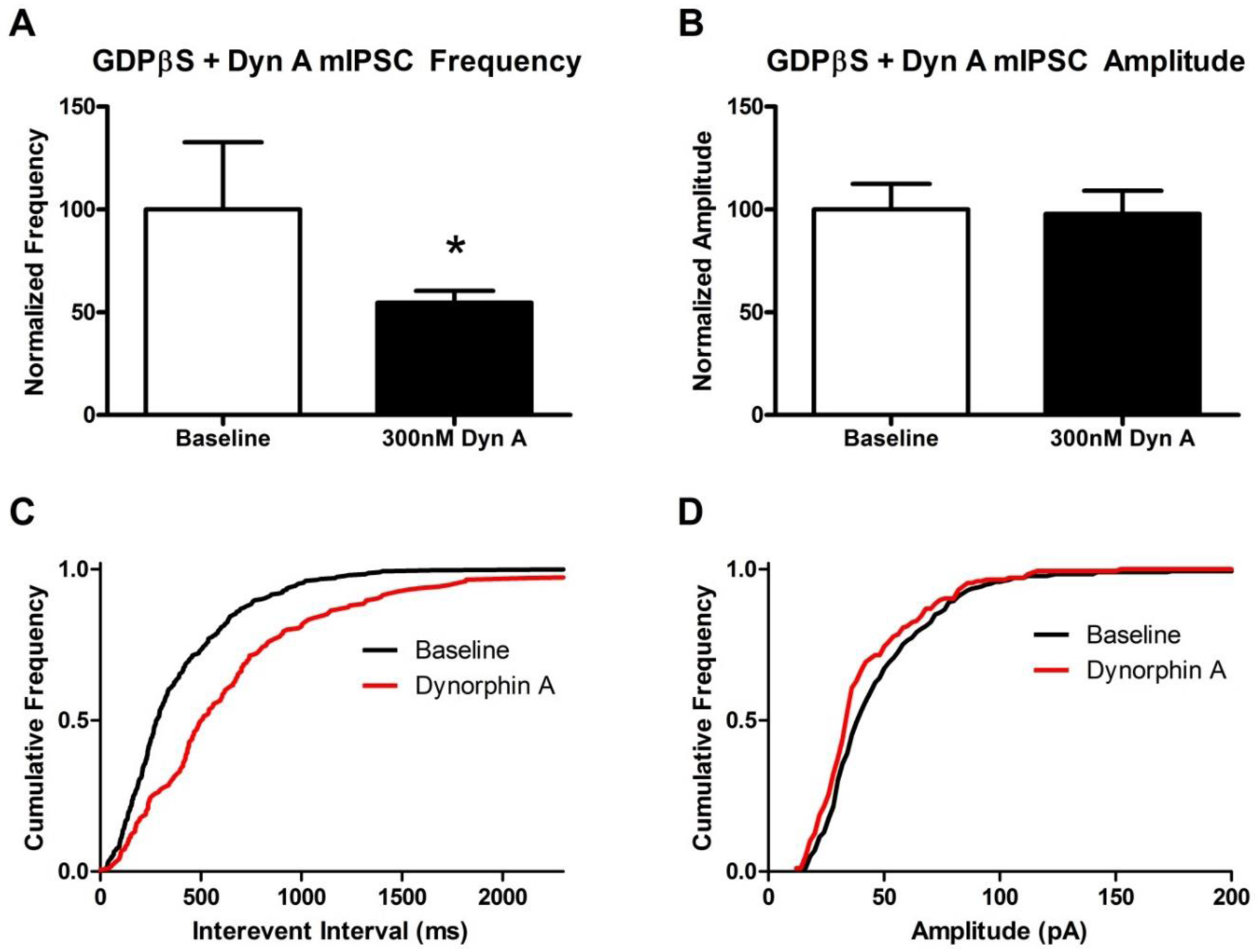
The postsynaptic GPCR inhibitor, GDPβs (4mM), defused into the recorded vlPAG DA neuron did not block dynorphin A from attenuating mIPSC frequency, suggesting a presynaptic mechanism. (A) Inhibition of postsynaptic GPCR did not block dynorphin A from attenuating mIPSC frequency. (B) Dynorphin A had no effects on mIPSC amplitude in the presence of GDPβs. (C) GDPβs did not block dynorphin A’s ability to shift the mIPSC cumulative frequency towards longer interevent intervals. (D) Dynorphin A had no effect on mIPSC amplitude cumulative probability

### Dynorphin effects on GABA is not mediated through MAP kinase signaling

To identify the downstream signaling through which KOR modulate GABAergic transmission, we examined the role of MAP kinases ERK1/2 and p38. Brain slices were incubated in either a selective MEK inhibitor (SL327, 10 µM, n=6) or p38 inhibitor (SB203580, 20 µM, n=5) for 40 min before and during dynorphin wash-on. In the presence of the MEK inhibitor SL327, dynorphin A significantly decreased mIPSC frequency (67.2±10.6% of baseline, p=0.04, Fig3A), but not amplitude (97.1±4.3% of baseline); In the presence of the p38 inhibitor SB203580, dynorphin A significantly decreased mIPSC frequency (46.6±9.9% of baseline, p<0.05, Fig3B) and amplitude (83.2±3.6% of baseline, p=0.02, Fig3B). Neither SL327 nor p38 altered the dynorphin-induced attenuation of GABAergic input onto vlPAG DA neurons, suggesting this observation was not mediated through MAP kinase signaling.

**Figure 3.**
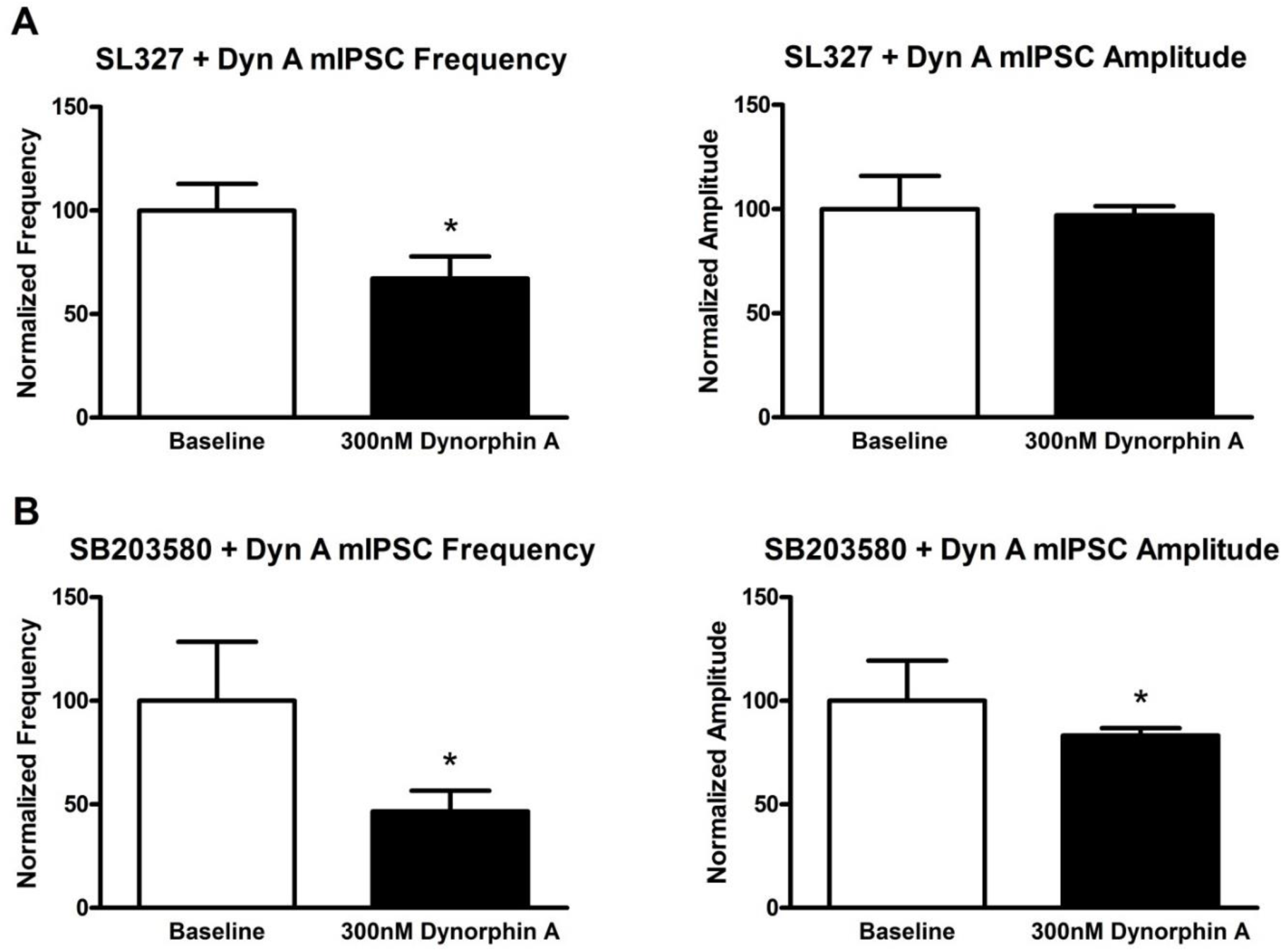
The ERK1/2 inhibitor SL327 (10 µM) and the p38 inhibitor SB203580 (20 µM) did not block dynorphin A from attenuating mIPSC frequency. (A) Dynorphin A significantly decreased mIPSC frequency in the presence of SL327. (B) Dynorphin A had no effects on mIPSC amplitude in the presence of SL327. (C) Dynorphin A significantly decreased mIPSC frequency in the presence of SB203580. (D) Dynorphin A significantly decreased mIPSC amplitude in the presence of SB203580

### Dynorphin effects on GABA are not mediated through calcium and potassium ion channel conductance

Agonist-induced dissociation of the βγ subunit from the GPCR can directly influence the conductance of ion channels. Thus, we investigated the roles of calcium and potassium channels in dynorphin modulation of GABA-mediated IPSC. To eliminate the role of calcium channels, we incubated the slices in calcium-free ACSF and the selective calcium chelators BAPTA-AM (50µM) and EGTA (100µM) for 1-2 hours before recording, and continued to record from the slice in calcium-free ACSF in the presence of just EGTA, or EGTA plus 4-AP (100µM) to block potassium channels. In calcium-free experiments, dynorphin A significantly decreased mIPSC frequency (71.9±8.6% of baseline, p=0.02, n=6, Fig4A), but not amplitude (102.3±9.6% of baseline, Fig4A). In calcium-free experiments where potassium channels were blocked with 4-AP, dynorphin A still caused a significant decrease in mIPSC frequency (77.8±8.0% of baseline, p<0.05, n=8, Fig4B), but not amplitude (107.5±5.7% of baseline, Fig4B). The persistence of KOR-mediated attenuation of IPSCs in the vlPAG dopamine neurons was not attribuated by the change in conductance of these ion channels. We explored the role of KOR βγ subunits beyond direct influence on ion channels by incubating the slices in gallein (100µM), an inhibitor of G protein βγ subunit-dependent signaling. The previously observed KOR activation-induced reduction of GABA transmission was blocked in the presence of gallein (n=6), with no significant changes in mIPSC frequency (107.1±7.1% of baseline, Fig4C) or amplitude (102.8±7.4% of baseline, Fig4C). Together these data suggest that the effects of KOR on GABAergic transmission were mediated via βγ subunit signaling, but not through the change in conductance of ion channels.

**Figure 4.**
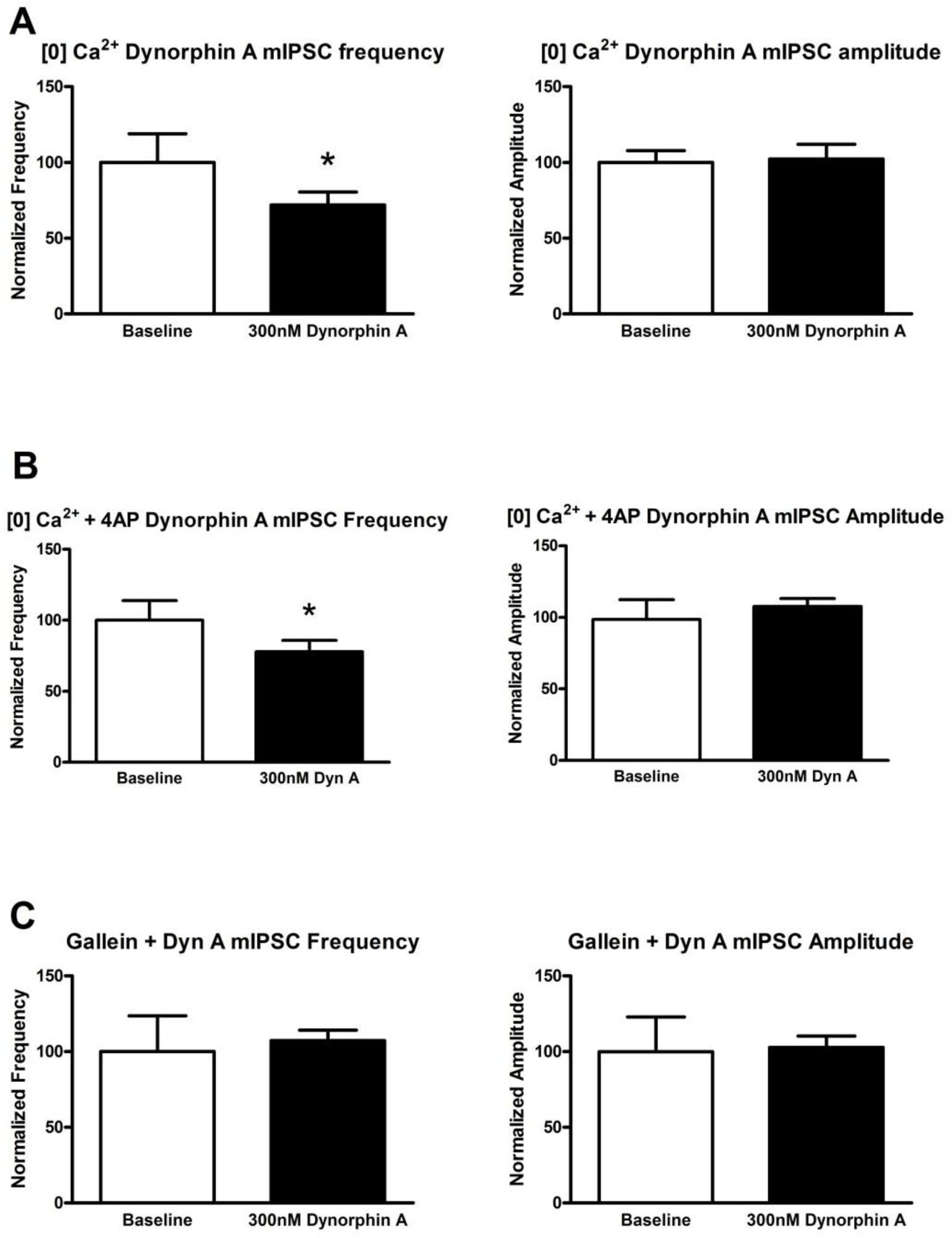
KOR inhibition does not require calcium and potassium ion channels. The GPCR βγ subunit inhibitor blocked dynorphin A from attenuating mIPSC frequency. (A) KOR/dynorphin A inhibition persisted even when all Ca^2+^ was removed by incubating slices in 0mM Ca^2+^/4mM Mg^2+^, 100µM EGTA with 50µM BAPTA-AM. (B) KOR/dynorphin A inhibition persisted even when all Ca^2+^ was removed and K^+^ channels are blocked with 4AP (100µM). (C) Gallein (100µM) prevented dynorphin A from inhibiting mIPSC frequency. mIPSC amplitude remained unaffected

**Figure 5.**
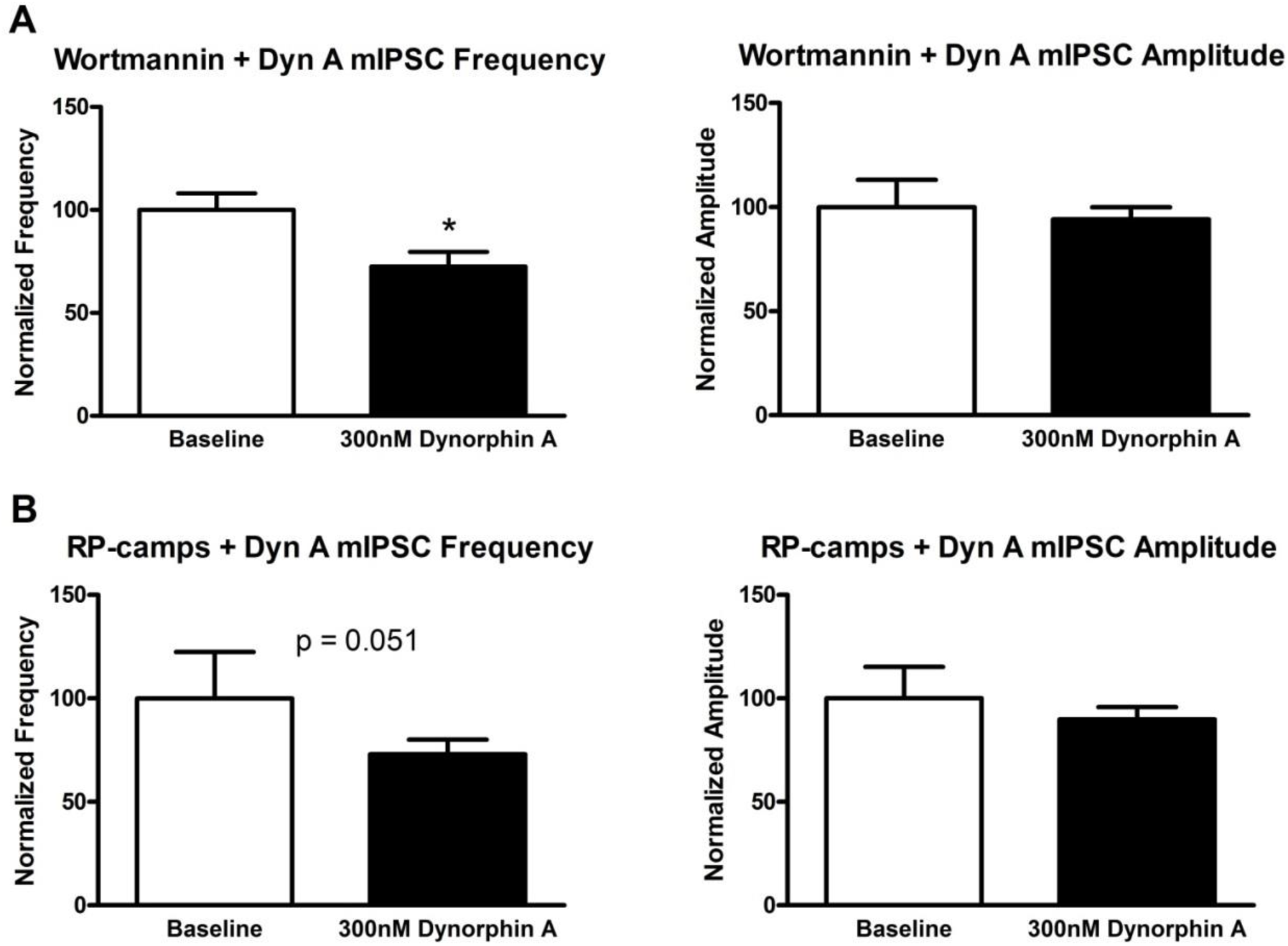
The PI_3_ kinase inhibitor wortmannin (1µM) and the PKA inhibitor RP-camps (10µM) did not block dynorphin A from attenuating mIPSC frequency. (A) Dynorphin A significantly decreased mIPSC frequency but had no effect on amplitude in the presence of wortmannin. (B) Dynorphin A decreased

Having established the possibility of βγ-mediated decrease in GABA function by dynorphin, we continued to delve into other downstream signaling of KOR activation -- PI3 kinase and PKA signaling. We incubated the slices in the PI3 kinase inhibitor wortmannin (1µM), or the PKA inhibitor RP-camps (10µM). Wortmannin did not block the previously observed decrease in GABA transmission frequency (72.4±7.1% of baseline, Data not shown). In addition, RP-camps also did not block the KOR-induced decrease in GABA frequency (73.0±7.0% of baseline, n=5, p=0.051, Data not shown) and there were no effects on amplitude. These data suggest that PI3 kinase and PKA do not mediate KOR activation-induced decrease in GABA function in the vlPAG.

## Discussion

vlPAG dopamine neurons have been implicated in a variety of emotional behaviors, in particular the regulation of pain. This study focused on determining the effects of kappa opioid receptor-activation on the inhibitory synaptic transmission onto vlPAG dopamine neurons, in addition to the downstream signaling mechanisms through which the effects take place.

We first investigated the effects of KOR activation on GABA-mediated mini IPSC and found that KOR activation by its endogenous ligand dynorphin A attenuated inhibitory transmission frequency but not amplitude, suggesting a presynaptic mechanism. This KOR effect is consistent with previous studies in the BNST [12], and was abolished in the presence of a KOR antagonist nor-BNI in both regions, suggesting KOR-selective mediation. Although all vlPAG DA neurons recorded in this study showed a decrease in GABA-mediated IPSCs, studies in the VTA demonstrated that KOR attenuates inhibitory transmission onto dopamine neurons in a projection target-dependent manner [25,26]. Recent evidence showed that GABA afferent region could also play a role in the differential modulation of IPSC by KOR activation [27]. Interestingly, KOR antagonist nor-BNI alone had no effects on spontaneous inhibitory transmission, implying no tonic constitutive kappa modulation on vlPAG DA neurons. In order to verify that the attenuation on GABAergic transmission was indeed mediated by presynaptic signaling of KOR, and not postsynaptic, we blocked GPCR signaling in the recorded postsynaptic cell. The blockade of postsynaptic GPCR was achieved by the addition of GDPβs in the patching pipette, filling the recorded cell by diffusion. GDPβs is a non-hydrolyzable GDP analogue; it blocks postsynaptic GPCR signaling by preventing the exchange of GDP to GTP, thus preventing G protein activation. We found that the reduction in mIPSCs caused by dynohpin A persisted in the presence of postsynaptic GDPβs, suggesting that postsynaptic KORs were not involved in this observation. Using GDPβs to block postsynaptic GPCRs has been shown effective in the Kash lab where postsynaptic NPY Y1 receptor agonist effects were blocked in its presence. The above evidence demonstrated KOR mediates inhibitory inputs onto vlPAG DA neurons presynaptically, which is not surprising as KOR agonists have been shown to inhibit presynaptic GABA release in other brain regions, such as the BNST [12], and in the hypothalamus [28].

We next disected downstream signaling cascade that could mediate the KOR activation-induced attenuation of IPSCs in vlPAG DA neurons. Previous studies in the Kash lab demonstrated in the BNST that KOR-induced decrease in GABAergic current was mediated presynaptically via the classic MAP kinase ERK1/2 pathway. In the BNST, the blockade of ERK1/2 signaling by inhibitors U0126 and SL327 both occluded the KOR effects on GABAergic transmission that was previously seen. However, current investigation in the vlPAG suggested site-specific actions of KOR activation-triggered ERK1/2 signaling in the inhibition of presynaptic GABA as inhibition of ERK1/2 signaling via SL327 did not occlude dynorphin-induced reduction in GABAergic transmission. In our previous work in the BNST, we found that in the presence of the p38 MAP kinase inhibitior, SB203580, KOR inhibition of glutamate release was blocked. However, in the vlPAG, KOR inhibition of GABAergic transmission persisted with p38 inhibitor onboard. Curiously, both mIPSC frequency and amplitude decreased upon application of the KOR agonist dynorphin A, implying perhaps SB203580 modulated postsynaptic properties that makes vlPAG dopamine neurons sensitive to dynorphin A.

We continued to examine ion channels that are directly regulated by the KOR βγ subunit. Opioids depress transmission at many central synapses [29]; many studies have implicated an kappa opioid-mediated inhibition of presynaptic calcium channels [30,31] or the activation of presynaptic potassium channels [32,33]. However, other studies have ruled out the involvement of either channel [26,34,35]. In our experiments, it is noteworthy that under zero calcium recording condition, the frequency of mIPSC more than halved in comparison to recordings under normal calcium concentration (refer to Table 1). The resulting marked and persistent attenuation in mIPSC frequency while having no change in amplitude upon application of dynorphin A is even more convincing under such low frequency baseline. Our data demonstrated that in the vlPAG dopamine neurons, neither of these ion channels contributed to the presynaptic inhibition of GABAergic release. However, inhibition of G protein βγ subunit-dependent signaling successfully prevented dynorphin-induced decrease in mIPSC frequency, suggesting that other actions of the βγ subunit can contribute to this effect. Because gallein has been shown to not only inhibit βγ subunit, but also activate PI_3_ kinase activity [36], we sought to clarify the role of PI_3_ kinase with wortmannin, a PI_3_ kinase inhibitor. Our results suggested that KOR-induced reduction of presynaptic inhibition lies outside of the actions of PI_3_ kinase. Together, we have confirmed that KOR-activation reduces inhibitory input via a presynaptic mechanism. In addition, with the removal of calcium, the effect persists, indicating that this inhibition occurs downstream of calcium entry and is calcium-independent. These results raised the possibility that dynorphin A could be activating KORs and directly affect the presynaptic release machinery in the GABAergic inputs onto the vlPAG dopamine neurons. This data is similar to studies on KOR presynaptic inhibition of glutamatergic inputs in the hypothalamus [35]. Although we did not identify the molecular target through which dynorphin inhibits presynaptic GABA, our results are consistent with studies proposing direct modulation of the exocytosis release machinery by the βγ subunit of the Gi/o-coupled GPCR [37,38]. We continued to rule out the possibility of the actions of phosphorylated KOR, inhibiting PKA and the generation of cAMP. We found a close to significant (p=0.051) decrease in mIPSC frequency upon the application of dynorphin A, consistent with other results that suggest KOR does not inhibit through that actions of PKA.

**Table 1.** Frequencies of IPSCs under various drug treatment

| Drug | Treatment | mIPSC frequency (±SEM) |
| --- | --- | --- |
| Dynorphon A | ----- | $1.4 \pm 0.4$ (n=5) |
| | 4AP | $1.1 \pm 0.4$ (n=7) |
| | [0] $\text{Ca}^{2+}$ | $0.4 \pm 0.1$ (n=6) |
| | [0] $\text{Ca}^{2+}$ + 4AP | $0.6 \pm 0.1$ (n=8) |
| | GDPβs | $1.3 \pm 0.4$ (n=6) |
| | Gallein | $1.6 \pm 0.4$ (n=6) |
| | norBNI | $1.1 \pm 0.2$ (n=5) |
| | RP-camps | $1.4 \pm 0.3$ (n=5) |
| | SB203580 | $1.7 \pm 0.5$ (n=5) |
| | SL327 | $1.1 \pm 0.1$ (n=6) |
| | Wortmannin | $1.0 \pm 0.1$ (n=6) |
| nor-BNI | ----- | $1.0 \pm 0.2$ (sIPSC) (n=6) |

In this current study, we found that KOR activation presynaptically attenuates inhibitory inputs potentially via the modulation of release machinery. However, KOR influences on excitatory inputs of vlPAG dopamine neurons are unclear. Some studies have shown that KOR does not modulate EPSCs onto other dopamine-rich regions, such as the VTA [39] while others have shown the opposite [40]. Over all cell excitability in the VTA has been shown modulated in a projection-specific manner [21] where only mPFC-projecting VTA dopamine neurons are hyperpolarized by KOR activation, but not NAc-projecting dopamine neurons. It is currently unclear how KOR modulation of vlPAG DA neurons alters behavior. Given the prominent role that the vlPAG plays in pain and negative affect processing, as well as the correlation of KOR functions and emotional behaviors, it is tempting to speculate that KOR actions on this circuit are involved in these processes. Under the hypothesis that KOR presynaptically inhibits GABAergic inputs, disinhibiting the vlPAG dopamine neurons to potentially modulate projection areas and related behaviors, further elucidation is needed regarding how the KOR modulates the glutamatergic inputs, as well as the overall effect on activity of the vlPAG DA neurons. Future studies utilizing optogenetics to probe pathway-defined plasticity, as well as applying designer receptors exclusively activated by designer drugs (DREADD) to characterize pathway and cell type-specific modulation will likely shed light on this exciting possibility.

